# Burnout in nurses and biomarkers of stress, inflammation and neuroplasticity

**DOI:** 10.1101/2025.03.11.642730

**Authors:** Bengt B. Arnetz, Jackeline Iseler, Michelle Pena, Nicolina Evola, John vanSchagen, John S. Beck, Scott E. Counts, Eamonn Arble, Judith E. Arnetz

## Abstract

Burnout is an occupational challenge to the health, performance, and retention of healthcare personnel. The objective of this cross-sectional study was to further our understanding of the association between burnout, work, coping, and cognitive impairment as it relates to neuroendocrine, inflammatory, and neuroplastic disease mechanisms. One hundred hospital- based registered nurses responded to a validated survey addressing employment and work characteristics, coping, and cognitive impairment, and a one-item, burnout scale. In addition, they all provided blood samples. Nineteen percent of the nurses reported symptoms of evolving burnout and an additional 12% reported established burnout. Severity of burnout was inversely associated with self-rated energy (p<.001), ability to concentrate (p<.001), and positively associated with stressed at work (p<.001), but not with workplace cognitive impairment. The anti-inflammatory and pro-energetic biomarker interleukin-10 was elevated in respondents in the combined two highest burnout categories (mean 2.81, S.E.M. 0.26 pg/mL) vs. a median of 2.09 pg/mL in the no-burnout category (p<.02). When biomarkers in blood were regressed on severity of burnout, concentration of the anabolic hormone dehydroepiandrosterone-sulfate (standardized beta -.73, p=.007) and the neuronal strain biomarker neurofilament light chain (-.79, p=.01) inversely predicted burnout. In contrast, the ability to cope with a tough situation at work was positively associated with burnout (.75, p=.02). The study not only confirms the association between burnout and self-reported individual and work-related adverse outcomes but, importantly, burnout-relevant neuroendocrine, inflammatory, and neuronal biomarkers. Nurses suffering from burnout might exhibit dysfunctional coping resulting in decreased recognition of low energy, which accelerates the burnout process. It is proposed that assessment of biological disease mechanisms should play a larger role in both scholarly and clinical burnout work.

## Introduction

The term “burnout” was first mentioned in 1969 [1]. However, the term was popularized by psychotherapist Herbert Freudenberger, who in 1974 reported how he and his colleagues personally experienced exhaustion, cynicism, and reduced work productivity. These three components are now recognized as the basic core of the burnout construct [2]. Since then, over 140 definitions of the phenomenon have been published [3]. Consequently, there are many measurement instruments, of which the Maslach Burnout Inventory (MBI) is the most well- known (4).

The current understanding of possible contributing causes to burnout includes a complex interplay between personal, professional, leadership, and organizational factors [5–11]. Sustained and unresolved workplace stress, poor fit between job demands and personal resources, ineffective leadership, and organizational inefficiency are commonly cited risk factors [8–10, 12–15]. However, as noted in an extensive review of the burnout literature by Rand Corporation, “Despite the general agreement over the components of burnout, the causes and consequences of burnout have not yet been defined.” [16].

In 2019, the World Health Organization (WHO) defined burnout as “a syndrome conceptualized as resulting from chronic workplace stress that has not been successfully managed.” [17, 18]. When burnout was included in the International Classification of Diseases (ICD)-11, WHO classified burnout as an occupational phenomenon, not a medical condition. The lack of a gold standard for defining and measuring burnout and the increasing challenge of getting busy professionals to fill out extensive surveys have popularized single-item burnout measures based on the person’s feelings of burnout [9]. There are also biologically validated single-item and 3-item instruments that assess self-rated energy and self-rated fatigue, respectively, constructs closely associated with burnout [20–22]. Developing a better understanding of the pathophysiological mechanisms behind burnout is critical. To effectively diagnose and treat burnout, we need to better understand causal risk factors vs. confounders, that is, non-casually related factors associated both with risk factors and burnout. Moreover, valid diagnosis and treatment of burnout require an in-depth understanding of causal burnout disease mechanisms [3]. Such knowledge will contribute to improved quality of life in the workforce.

Addressing burnout might also improve the efficiency and quality of patient care since burnout is associated with decreased productivity, near misses, and adverse patient outcomes [9, 23, 24].

The conservation of resources (COR) model suggests that burnout is the result of the depletion of resources required to functionally manage stressors, resulting in dysfunctional coping [25, 26].

In the following, we discuss burnout from a neurophysiological perspective.

Neurocognitive symptoms of burnout suggest dysfunction in the limbic system, including the hypothalamus, hippocampus, amygdala, and septal nuclei [4]. Over a decade ago, Jovanovic et al. reported a functional disconnection between the amygdala and the anterior cingulate cortex (ACC) and reduced 5-HT_1A_ receptor binding potential in the ACC, the insular cortex, and the hippocampus in chronically stressed vs non-stressed subjects [27]. Subsequently, Blix et al. published a MRI study of highly stressed patients, that had been diagnosed as having had a “reaction to severe stress and adjustment disorder” according to the International Classification of Disease (ICD-10, F42) vs healthy controls [28]. The authors reported reduced grey matter volumes of the anterior cingulate cortex and the dorsolateral prefrontal cortex in highly stressed vs control participants. Moreover, the volumes of the caudate and putamen were reduced in highly stressed participants and the volume correlated inversely to the degree of perceived stress. The stress-adaptive, catabolic, and neurotoxic hormone cortisol plays a key role in the regulation and functioning of the limbic system [29–31]. The hypothalamic-pituitary-adreno-cortical axis has been reported to be up and down-regulated in burnout [29, 32, 33]. Impaired neurogenesis, decreased levels of the neuroprotective and anabolic hormone dehydroepiandrosterone-sulfate (DHEA-s), and increased concentration of biomarkers of systemic inflammation, such as, interleukin-6 and tumor necrosis factor-alpha (TNF-α), might contribute to neurocognitive impairment symptoms associated with burnout [4, 34, 35]. Furthermore, increased C-reactive protein (CRP) concentration has been related to decreased cognitive function [36].

Fatigue, or low energy syndrome, is an important and quality-of-life limiting component of burnout related to reduced levels of interleukin-10, a key anti-inflammatory component of the body’s inflammatory regulatory system [20, 22, 37]. As discussed above, burnout and work stress impair the ability to concentrate at work and might contribute to work-related cognitive impairment in providers [24, 28, 38]). Impairment in concentration ability is hypothesized to contribute to lapses in memories, workplace cognitive impairment and failure, that is, errors in cognitive processing and decision-making [39, 40]. Yet while the brain plays a central role in burnout, no studies to date have simultaneously examined both neurobiological and workplace risk factors for burnout [41, 42]. It is well-recognized that exposure to intense psychological trauma is associated with functional and epigenetic changes reflective in peripheral brain markers in blood [43, 44]. Still, even the extensive Rand Corporation review of the burnout literature lacked a section dedicated to the physiology of burnout [16]. There is a void of neurobiological studies exploring the link between work-related cognitive dysfunction and neurogenesis. However, a few burnout studies have applied a neurotrophic model [28, 45]. Of special interest is the role of impaired synthesis of neurotrophins, such as brain-derived neurotrophic factor (BDNF) and nerve growth factor (NGF), as well as vascular endothelial growth factor (VEGF) and epidermal growth factor (EGF), in burnout [4, 46, 47]. These biomarkers are part of the brain’s adaptive responses to stress, promoting neuronal restoration and growth and countering programmed neuronal cell death. Department-level workplace stress, including bullying [48], a burnout risk factor, has been associated with decreased concentration of the neuroprotective hormone DHEA-s in nurses. Moreover, it has been related to adverse patient outcomes at the department level [34]. Since burnout is not a medical diagnosis, different criteria have been used to define groups as burned out, resulting in difficulties comparing results (3). Most studies focus on the emotional exhaustion component of burnout when defining participants as burned out [47]. We hypothesize that sustained stress and burnout result in decreased levels of neurotrophic factors and increased levels of neuronal strain. Indeed, burnout symptoms in healthy subjects have been associated with lower levels of BDNF [49]. Self-rated mental stress has been reported to be inversely related to levels of BDNF [50]. These findings have clinical implications since workplace stressors, antecedents to burnout, are associated with increased risk for central line-associated bloodstream infection (sepsis) in patients, possibly due to workplace cognitive impairment [38, 48].

The overall objective of the current study was to determine the possible association between burnout category severity, workplace stress and cognitive impairment, inflammation, neuroendocrine and neuroplastic profiles in hospital-based nurses with a special focus on the prefrontal area and the limbic systems. Fig 1 depicts the brain neurobiological model informing the study design and analysis.

**Fig 1.**
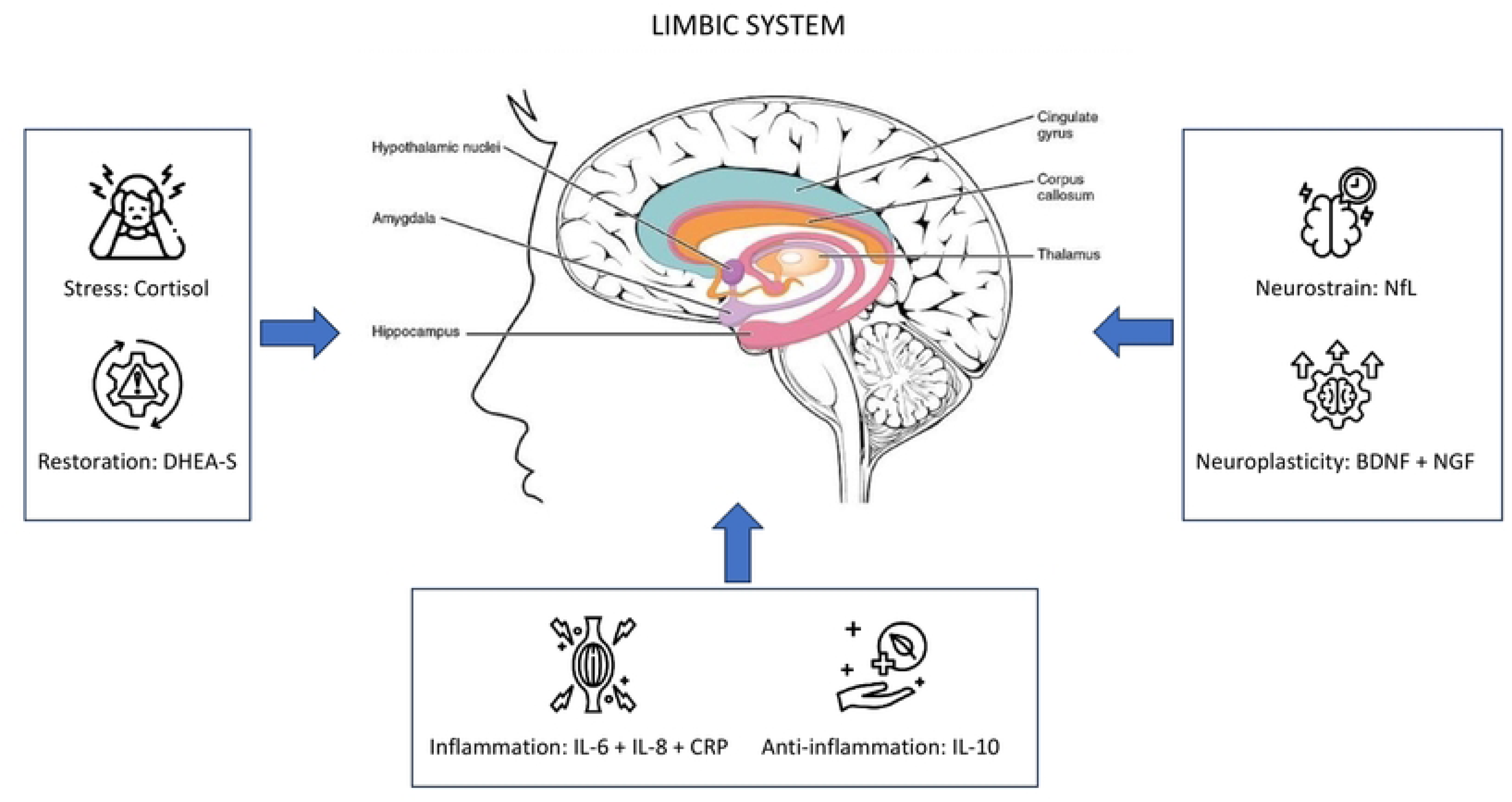
Neurophysiological burnout model. Theoretical model that informed the study design and analysis depicting the relationship between pro- and anti-neuronal growth factors and their association to the limbic and associated systems hypothesized to be causally related to burnout. Legend. IL = interleukin. DHEA-s = Dehydroepiandrosterone-sulfate. CRP = C-reactive protein. NfL= Neurofilament Light. BDNF = brain- derived neurotrophic factor. NGF = Nerve growth factor.

The current study was guided by the following three inter-related hypotheses:

1. Burnout <u>categories</u> are associated with decreased energy, decreased ability to concentrate, and increased workplace cognitive impairment.
2. Burnout <u>categories</u> are associated with stress-adaptive neuroendocrine, pro-, and anti-inflammatory patterns as well as neuroplastic and neuronal strain.
3. Lower energy, lower ability to cope with hardship at work, and stress-adaptive neuroendocrine, inflammatory, neuroplastic, and neuronal strain systems predict burnout severity.

## Methods

This cross-sectional study took place in 2024 at a community teaching hospital in West Michigan, U.S., part of a large national hospital system. The hospital has 5126 employees, of which 1,166 are direct care registered nurses, and an additional 305 “other” registered nurses in management and quality improvement roles. The overall goal of the main study was to assess the psychometric properties and practical feasibility of a new instrument to assess thriving in healthcare personnel. The current paper focuses on the neurobiology of burnout in the 100 nurse participants who responded to a self-report survey and provided blood samples.

## Questionnaire

The Thriving in Nursing Questionnaire (THINQ) is based on previously validated self- reported items and scales discussed below [51, 52]. The THINQ survey contained 29 questions, of which 7 were demographic items measuring hospital unit, work shift, years of nursing experience, age, gender, Hispanic/Latino origin, and race/ethnicity [53].

The THINQ survey also included self-reported questions on work, health, well-being, and burnout, as described below. To reduce the burden on participants, visual analog scales (VAS) were used rather than multi-item scales [53]. Items were selected from the validated Arnetz and Hasson Stress Questionnaire, an instrument comprised of seven VAS items based on the Quality, Work, Competence questionnaire measuring the psychosocial work environment (15, 51, 54). Research has found that these VAS items can be used instead of Likert-type items and have sound psychometric properties [51, 55].

The following questionnaire items were used in the current study:

Self-rated health: A single item, “how is your overall health right now?” was a VAS item ranging from 0 (very poor) to 10 (very good). Self-rated health was included as a quality check to ensure the sample based on the 100 nurses confirmed published results and hypothesized associations between self-rated health and a person’s future health projection and health-related biomarkers [56, 57].

The following VAS items were used to test the hypotheses presented above.

“How is your energy right now? “[20, 51] scored from 0 (very poor) to 10 (very good).

How is your ability to concentrate right now?” [51, 54] scored from 0 (very poor) to 10 (very good).

The ability to cope with workplace challenges was assessed by the item: “I am able to handle a tough situation at work,” rated on a VAS scale from 0 (completely disagree) to 10 (completely agree).

The current study focused on cognitive impairment at work, defined as lapses in perception, memory, and actions during tasks that an individual would generally be able to carry out (58). Therefore, the item “I have lapses in memory at work” was selected from the Attention subscale of the Workplace Cognitive Failure scale [59, 60]).

Burnout was assessed using a single question developed by Rohland et al. [61]). The participant responded to the question: “Overall, based on your definition of burnout, how would you rate your level of burnout?” The five possible response alternatives were:

1. “I enjoy my work. I have no symptoms of burnout.” Referred to as (No-burnout).
2. “Occasionally, I am under stress, and I don’t always have as much energy as I once did, but I don’t feel burned out.” (Stressed)
3. “I am definitely burning out and have one or more symptoms of burnout, such as physical and emotional exhaustion.” (Evolving)
4. “The symptoms of burnout that I’m experiencing won’t go away. I think about frustration at work a lot.” (Burnout)
5. “I feel completely burned out and often wonder if I can go on. I am at the point where I may need some changes or may need to seek some sort of help.” (Burnout)

Based on published validation work, the single burnout question shows a correlation of 0.5, p<.0001 (using paired ANOVA and simple Pearson correlation coefficient) with the emotional exhaustion subscale of the Maslach Burnout Inventory [61]. The single-item scale is widely used in healthcare settings [62].

The final question in the survey asked for 100 nurses willing to provide blood on a single occasion. Those who agreed to provide a blood sample were asked to provide their contact information so they could schedule the blood draw.

### Blood sampling

The first 100 nurses to volunteer to provide blood were scheduled for individual blood draws in the hospital’s clinical laboratory. Following additional information about the study and analysis their blood would be subject to, consent was secured, and blood was sampled. From each nurse, a total of 20 mL of blood was collected using 1 x 10 mL red-top glass tube that was untreated and 1 x 10 mL purple-top glass tube treated with the anti-coagulant ethylenediamine- tetraacetic acid (EDTA). The red top tube was left to clot for 30 minutes and then centrifuged at 4 degrees Centigrade for 15 minutes at 3,500 rounds per minute (rpm). The resulting supernatant represented serum used for the analysis of specified biomarkers below. The purple top tube was centrifuged at 4 degrees Centigrade for 15 minutes at 3,500 rpm. The supernatant represented plasma used for subsequent analysis of below specified biomarkers.

The following biomarkers were chosen to reflect the various neurobiological systems of interest: 1. serum concentrations of cortisol and dehydroepiandrosterone-sulfate to gauge the body’s balance between stress, or catabolic, and restorative, or anabolic, processes. 2. Serum concentrations of CRP, interleukin (IL)-6, and IL-8 to assess systemic inflammation. 3. Serum concentration of IL-10 to gauge pro-energetic and anti-inflammatory processes. Serum concentrations of Il-6, IL-8, and CRP were added up to create an *inflammatory index*. Neuronal processes were measured using the two major neurotrophins, plasma BDNF and NGF, and the concentration in plasma of neuronal strain indicator neurofilament light chain (NfL). A *neuroplasticity index* was created by adding concentrations of BDNF and NGF. Commercial, human-specific 96 well ELISAs were used to measure the <u>serum</u> analytes as follows: cortisol (ALPCO #11-CRLHU-E01, Salem, NH; sensitivity = 0.4 ug/mL), CRP (ALPCO #30-9710S; sensitivity = 0.124 ng/mL), DHEA-s (ALPCO #11-DHEHU-E01; 5 ng/mL), IL-6 (BioLegend #430507, San Diego, CA; 1.6 pg/mL), and IL-8 (Boster Bio #EK0413, Pleasanton, CA; <1 pg/mL), and IL-10 (BioLegend #430607, IL-10; 2 pg/mL). The following human-specific ELISA kits were used to measure the <u>plasma</u> analytes of BDNF (Boster Bio #EK0307; sensitivity <15 pg/mL) and NGF (AVIVA #OKEH00186, San Diego, CA; 15.6 pg/mL). <u>Plasma</u> NfL was measured using the SIMOA HD-1 platform (Quanterix #104073, Billerica, MA; sensitivity =1.38 pg/mL). The manufacturer’s instructions were followed for all assays. Duplicate samples were quantified using standard curves based on calibrators of known concentration; intra-assay % CVs ≤ 5.0 % for all assays.

### Study participants

Before distributing the survey, nurse leadership arranged several meetings with nurses to inform them about the purpose of the upcoming “Nurse Thriving and Well-Being Study.” Moreover, the hospital’s Nursing Scholarly Practice Council and the Nurse Practice Environment Council (NPEC) were informed and supportive of the study. All registered direct care nurses employed by the hospital (n=1,166) were eligible to participate in the study. Posters informing about the study were mounted in areas of the hospital frequented by nurses. The hospital’s nurse leadership sent an invitation by email to all eligible nurses. The email explained the purpose of the study.

Nurses were informed that all responses would be anonymous, and individual responses could not be linked to their names or emails. The recruitment period started March 26, 2024, and ended May 3, 2024. Nurses who agreed to participate accessed the electronic Qualtrics survey by scanning the QR code or using the weblink provided in the email. The first page of the survey explained that participation was voluntary. The final question in the survey asked for 100 nurses willing to provide blood samples on a single occasion. Those who agreed to provide a blood sample were asked to provide their contact information so they could schedule the blood draw. All blood samples were taken at an on-site lab at the hospital. Completing the survey constituted nurses’ consent to participate.

### Data Collection

The survey was open from late March to early May 2024. Over the five-week study period, nursing leadership posted weekly reminders on the hospital’s nursing social media page. Of the 1,166 surveys distributed, 252 were returned, representing a 21.6% response rate.

### Comparing respondents to the survey only with those that also provided blood

Regarding the 100 nurses that also provided blood samples, and thus part of this paper, 95.0% identified as female, 95.0% as Caucasian, 29.0% aged 25-34, 25.0% aged 35-44, and 38.1% 45 years and older. Fourt persons (4%) identified as American Indian/Alaska Native, Asian, or Black/African American. Among nurses that only responded to the survey, 89.5% identified as female, 93.9% as Caucasian, one nurses (1%) belonging to another race/ethnic group, and 7 (7%) declined to answer the question. Differences between groups in terms of gender distribution (F2=3.01, two-sided p-value=.22) and race/ethnicity (F_4_=3.01, p=.22) were not statistically significant. The age distribution among survey respondents only was: 32.7% aged 25-34, 24.6% aged 35-44, and 24.6% aged 45 year and older. The age distribution did not differ significantly between the two groups (F_6_ = 6.64, p=.36).

The majority (84.0% vs 78.8%) worked day shift (F_1_=.96, p=.33 between groups). Nurses providing blood had worked longer in the nursing profession (17.2% vs 23.9% had worked less than 5 years; 23.2% vs 34.5% 5-20 years; and 59.6% vs 42.6% had worked longer than 10 years, F_2_=6.87, two-sided p-value=.03.) Nurses who provided blood rated their energy higher (mean = 6.73 out of a max of 10 [S.D. = 1.93]) versus 6.22 [2.03] in nurses that only responded to the survey (t_243_ =243, p=.04.) There were no group differences regarding the ability to concentrate. Nor was there any significant difference in burnout self-assessment across the five different response alternatives between nurses who responded to the survey only and those who also provided blood.

### Comparing survey responders to the entire nurse population of the hospital

Based on gender (92.1% female) and ethnic identification (93.9% Caucasian) of respondents, the study population differed from the total population of hospital nurses: 87.7% female, Chi square=4.95, p<.05; 86.2% Caucasian, Chi square=21.5, p<.001).

### Statistical Analysis

Data was first quality-checked to ensure there were no apparent errors in coding. Mean and dispersion measures were checked to ensure that the statistical assumptions for chosen tests were met. Two-sided Student t-test and Chi-square statistics were used to compare continuous and discrete variables, respectively. A psycho-biological validation check was conducted before analyses for the specific hypotheses using self-rated health, a validated measure of a person’s future health perspective, and positively related to the anabolic hormone dehydroepiandro- sterone-sulfate (DHEA-s) [56, 57]. It was therefore hypothesized that self-rated health would be positively associated with the ability to concentrate and self-rated energy, but inversely associated with lapses in memory at work. Moreover, it was hypothesized that self-rated health would be positively associated with blood concentrations of DHEA-s and the two neurotrophins, but inversely associated with the stress hormone cortisol, the inflammatory index, and the neuronal strain biomarker NfL.

Linear regression analysis was used to test hypothesized predictive models. In analyses of burnout, Category 4 (“The symptoms of burnout that I’m experiencing won’t go away”) and 5 (“I feel completely burned out”) were combined into category 4 and named Burnout. Category 1 was considered no burnout; Category 2 categorized nurses who were under stress and experienced less energy, but did not feel burned out; and Category 3 categorized nurses with evolving symptoms of burnout. A one-sample test was used to compare the mean of specific variables in the combined two highest burnout categories versus the median of the no-burnout category. Statistical significance was set at a two-sided p-value of <.05.

## Ethical considerations

The study was approved by the Institutional Review Boards at Michigan State University (Study ID 00009623) and Trinity Health Grand Rapids Hospital (Study ID 24-0122-4).

## Results

Based on the single-item burnout measure, almost seven out of ten participants responded that “I enjoy my work. I have no symptoms of burnout” (Category 1, No burnout, 11%) or “Occasionally I am under stress, and I don’t always have as much energy as I once did, but I don’t feel burned out” (Category 2, Stressed, 58%). Nineteen percent reported clear symptoms of evolving burnout, and 12% reported symptoms reflective of established burnout (original category 4, 10%, and category 5, 2%). Categories 4 and 5 were combined into category 4, Burnout, representing 12% of all respondents.

As hypothesized, self-rated health was associated with the ability to concentrate (r = .63, p<.001), self-rated energy (r=0.65, p<.001), and inversely to lapses in memory (r=-.26, p<.001). Furthermore, self-rated health was inversely associated with the inflammatory index (r=-.28, p=.046) and counter to the hypothesis, positively with NfL (r=.24, p=.018). However, there was no association between self-rated health and neurotrophins or serum-cortisol.

### Hypothesis 1

Increased Burnout severity categories are associated with decreased energy, decreased ability to concentrate, and increased workplace cognitive impairment.

Table 1 shows that there is an inverse association between advancing burnout severity categories and self-rated energy and ability to concentrate.

**Table 1.** Self-rated health variables across the four burnout categories.

| Variable | Burnout<br>Category<sup>#</sup>(n) | Mean<br>(S.D.) | F<sub>df</sub> value | p-value | Range | Anchor<br>s |
| --- | --- | --- | --- | --- | --- | --- |
| How is your<br>Energy right<br>now? | 1 (11) | 8.37 (1.69) | F<sub>3</sub> 10.66 | <.001 | 0-10 | Very<br>low,<br>Very<br>high |
|  | 2 (58) | 7.06 (1.71) |  |  |  |  |
|  | 3 (19) | 6.19 (1.77) |  |  |  |  |
|  | 4 (12) | 4.68 (1.50) |  |  |  |  |
| Ability to<br>Concentration<br>right now? | 1 | 8.23 (2.01) | F<sub>3</sub> 8.31 | <.001 | 0-10 | Very<br>low,<br>Very<br>high |
|  | 2 | 7.37 (1.59) |  |  |  |  |
|  | 3 | 6.74 (1.73) |  |  |  |  |
|  | 4 | 5.03 (1.92) |  |  |  |  |
| Lapses in<br>Memory at<br>work | 1 | 3.57 (3.65) | F<sub>3</sub> .69 | 0.56 | 0-10 | Never,<br>Always |
|  | 2 | 3.73 (2.76) |  |  |  |  |
|  | 3 | 3.85 (3.09) |  |  |  |  |
|  | 4 | 5.06 (3.28) |  |  |  |  |
| I feel Stressed<br>at work | 1 | 2.42 (1.60) | F<sub>3</sub> 19.23 | <.001 | 0-10 | Never,<br>Always |
|  | 2 | 4.80 (2.13) |  |  |  |  |
|  | 3 | 7.21 (2.01) |  |  |  |  |
|  | 4 | 7.68 (1.87) |  |  |  |  |

<sup>#</sup> Burnout categories 4 and 5 have been merged into burnout category 4.
Independent samples, two-sided T-tests. SD=standard deviation

There was a positive association between burnout categories and being stressed at work. There was no association between burnout category and lapses in memory at work. A significant dose-response association (p<.001) was seen across the four burnout categories and the above ratings, except for lapses in memory at work. With increasing burnout severity category, self- rated energy and ability to concentrate decreased while feeling stressed at work increased

### Hypothesis 2

Burnout severity <u>categories</u> are associated with stress-adaptive neuroendocrine, pro-, and anti-inflammatory patterns as well as neuroplastic and neuronal strain. Contrary to the a’ priori hypothesis, there was no statistically significant association across the four re-categorized categories of burnout and blood markers reflecting the four neurobiological systems.

Since biomarkers across burnout categories were the same, Table 2 depicts results for the respective blood biomarkers for respondents in aggregate, not split across the four burnout categories.

**Table 2.** Mean values for neurobiological biomarkers for all participants in aggregate. Test statistics represent comparisons across the four categories of burnout.

| Variable | Mean (S.D.) | F<sub>df</sub> value | p-value<sup>s</sup> | 95%-C.I.<sup>#</sup> |
| --- | --- | --- | --- | --- |
| S<sup>#</sup>-Cortisol, ug/dL | 6.38 (3.88) | F<sub>3</sub> .91 | .44 | 2.73-13.11 |
| S-DHEA-s, ug/mL | 1.55 (.79) | F<sub>3</sub> .23 | .87 | .39-3.08 |
| S-CRP, ng/mL | 6159.38 (5919.09) | F<sub>3</sub> .94 | .43 | 169.70-20942.82 |
| S-IL-6, pg/mL | 3.28 (4.04) | F<sub>3</sub> 1.14 | .34 | .21-10.63 |
| S-IL-8, pg/mL | 1.06 (1.79) | F<sub>3</sub> .60 | .62 | .03-5.43 |
| S-IL-10, pg/mL | 3.92 (5.31) | F<sub>3</sub> 1.57 | .20 | 1.13-11.53 |
| P<sup>&</sup>-BDNF, pg/mL | 148.67 (92.79) | F<sub>3</sub> .58 | .63 | 48.80-353.50 |
| P-NGF, pg/mL | 4.57 (6.18) | F<sub>3</sub> .29 | .83 | .14-17.87 |
| P-NfL, pg/mL | 5.37 (2.72) | F<sub>3</sub> 2.54 | .06 | 2.30-11.00 |
| S-Inflammatory Index*, ng/mL | 6743.74 (6611.94) | F<sub>3</sub> 1.45 | .24 | 377.78-21347.84 |
| P-Neuroplasticity Index**, pg/mL | 193.99 (109.81) | F<sub>3</sub> 1.19 | .32 | 66.73-456.15 |

<sup>#</sup>Serum
<sup>&</sup>Plasma
<sup>s</sup>Student’s t-test, two-sided p-value
<sup>#</sup>95%-confidence interval
\* Sum of IL-6, IL8, and CRP
\*\* Sum of P-BDNF and P-NGF

As indicated in the table, the 95% confidence intervals for the inflammatory and neuroplasticity indexes are wide. This reflects the large variation in systemic inflammation and neuroplastic activity across individuals.

To determine whether the most extreme categories of burnout differed on key outcome measures, self-ratings, and blood biomarkers, the two combined burnout categories, re-classified as category 4 (Burnout) were compared with the median value for respondents in category 1, No- burnout, using one-sample, two-sided t-tests. Table 3 shows that participants in the re- categorized burnout category 4 scored significantly lower on energy and ability to concentrate and higher on stress at work compared to those in the non-burnout category 1. There were no group differences for lapses in memory at work. Serum concentration of IL-10 was significantly higher in burnout category 4 than in the no-burnout category. Moreover, the concentration of log transforming (Ln), due to non-normal distribution, was lower in the burnout category.

**Table 3.** One-sample T-test comparing burnout category 1 vs the combination of 4 and 5 (cat 4).

| Variable | No-burnout<br>(No-BO – Cat 1) | No-BO | Burnout<br>(cat 4 and 5<br>combined) | t<sub>df</sub> | p-value<sup>@</sup> |
| --- | --- | --- | --- | --- | --- |
|  | Mean (S.D.)<br>n = 11 | Test<br>value<br>Median<br>n = 11 | Mean (S.D.)<br>n = 12 |  |  |
| Energy | 8.37 (1.69) | 9.00 | 4.68 (1.50) | t<sub>11</sub> - 9.99 | <.001 |
| Concentration | 8.22 (2.00) | 9.00 | 5.03 (1.92) | t<sub>11</sub> - 7.19 | <.001 |
| Lapses in memory | 3.57 (3.65) | 2.50 | 5.06 (3.28) | t<sub>11</sub> 1.49 | <.001 |
| Stressed at work | 2.42 (1.60) | 2.00 | 7.68 (1.87) | t<sub>11</sub> 11.26 | <.001 |
| S<sup>#</sup>-Cortisol, ug/dL | 4.78 (1.23) | 4.44 | 7.36 (6.49) | t<sub>11</sub> 1.56 | .15 |
| S-DHEA-s, ug/mL | 1.64 (.90) | 1.43 | 1.64 (1.05) | t<sub>11</sub> .73 | .48 |
| S-CRP, ng/mL | 7013.00 (6445.93) | 3709.85 | 4597.00 (6110.24) | t<sub>11</sub> .50 | .63 |
| S-IL-6, pg/mL | 1.87 (1.43) | 1.74 | 1.82 (1.75) | t<sub>11</sub> .16 | .88 |
| S-IL-8, pg/mL | 0.56 (.48) | .47 | .56 (.55) | t<sub>7</sub> .46 | .66 |
| S-IL-10, pg/mL | 6.93 (14.02) | 2.09 | 2.82 (0.88) | t<sub>11</sub> 2.86 | .016 |
| P<sup>S</sup>-BDNF, pg/mL | 135.83 (56.18) | 128.30 | 147.82 (86.94) | t<sub>11</sub> .79 | .45 |
| P-NGF, pg/mL | 4.13 (3.83) | 3.11 | 3.57 (2.48) | t<sub>5</sub> .46 | .67 |
| P-NfL<sup>^</sup>, pg/mL | 7.01 (3.00) | 6.58<br>[ln 1.62] | 5.26 (2.18)<br>[ln 1.58 (.42)] | t<sub>11</sub> - 2.09<br>[t<sub>11</sub>-41] | .06<br><.001<sup>^</sup> |
| S-Inflammatory Index*, ng/mL | 6750.85 (7503.90) | 2687.88 | 2900.68 (3731.72) | t<sub>6</sub> .15 | .89 |
| S-Neuroplasticity<br>Index**, pg/mL | 173.06 (49.12) | 182.00 | 206.98 (86.80) | t<sub>5</sub> .71 | .51 |

There were no significant differences, using the median for burnout Category 1 (No- burnout) vs Cat 4 and 5 combined (Burnout) for Cortisol, DHEA, CRP, IL-6, IL-8, BDNF, NGF, the Inflammatory Index, or the Neuroplasticity Index.

### Hypothesis 3

lower energy, lower ability to cope with hardship at work, and stress-adaptive neuroendocrine, inflammatory, neuroplastic, and neuronal strain systems predict burnout severity.

Linear analysis was used to predict burnout category scores (dependent) using defined self-rated and blood biomarkers as independent or predictor variables. Table 4 display the statistically significant model. Self-rated energy, serum levels of DHEA-s, and plasma levels of NfL were inversely related to burnout scores. That is, decreasing levels of self-rated energy, DHEA-s, and NfL predicted higher burnout scores. The variable used to assess coping strategy at work – “I am able to handle a tough situation at work” was positively associated with burnout category. The participants’ typical work schedule (day vs nigh-time shift) did not predict burnout severity.

**Table 4.**
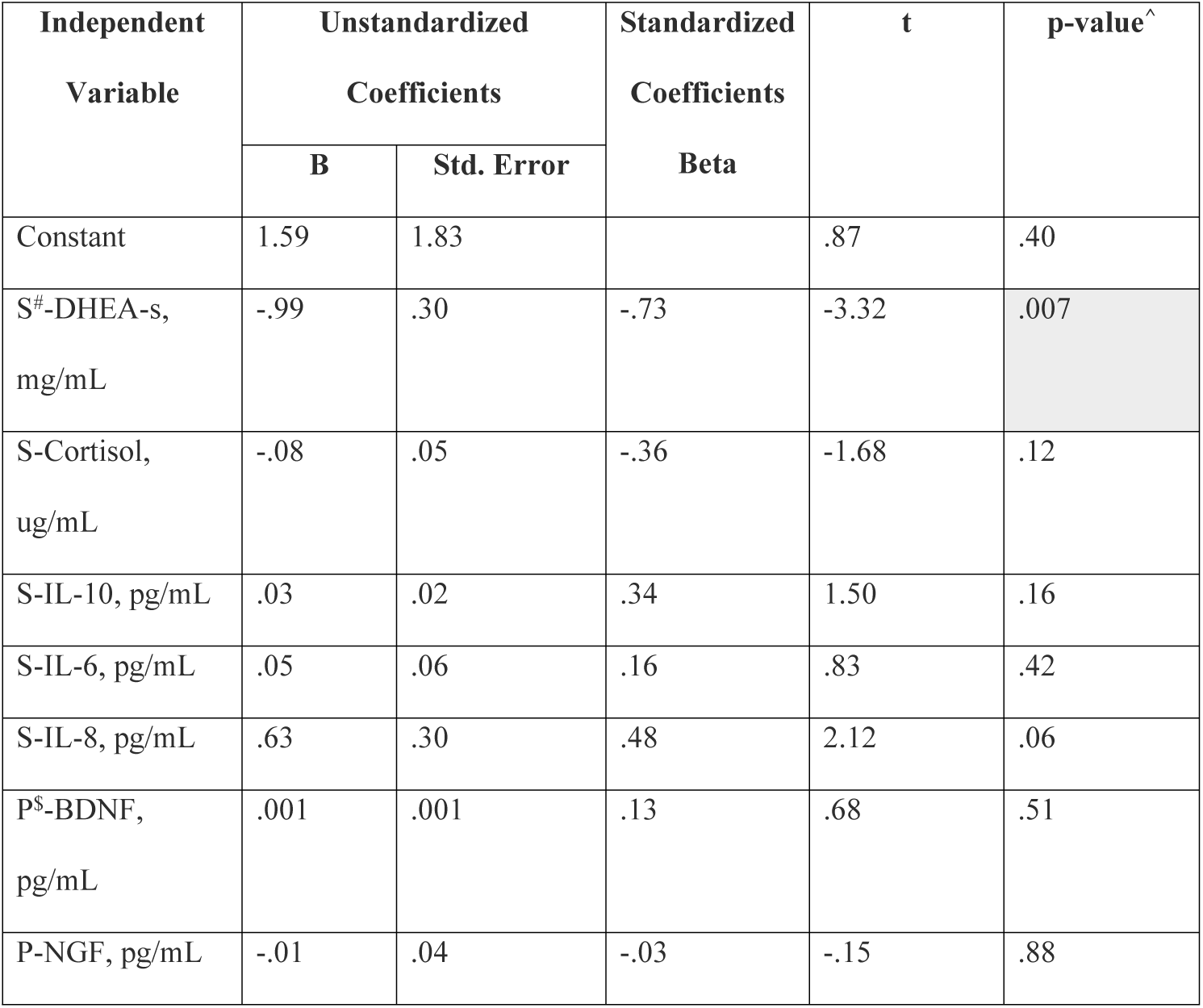
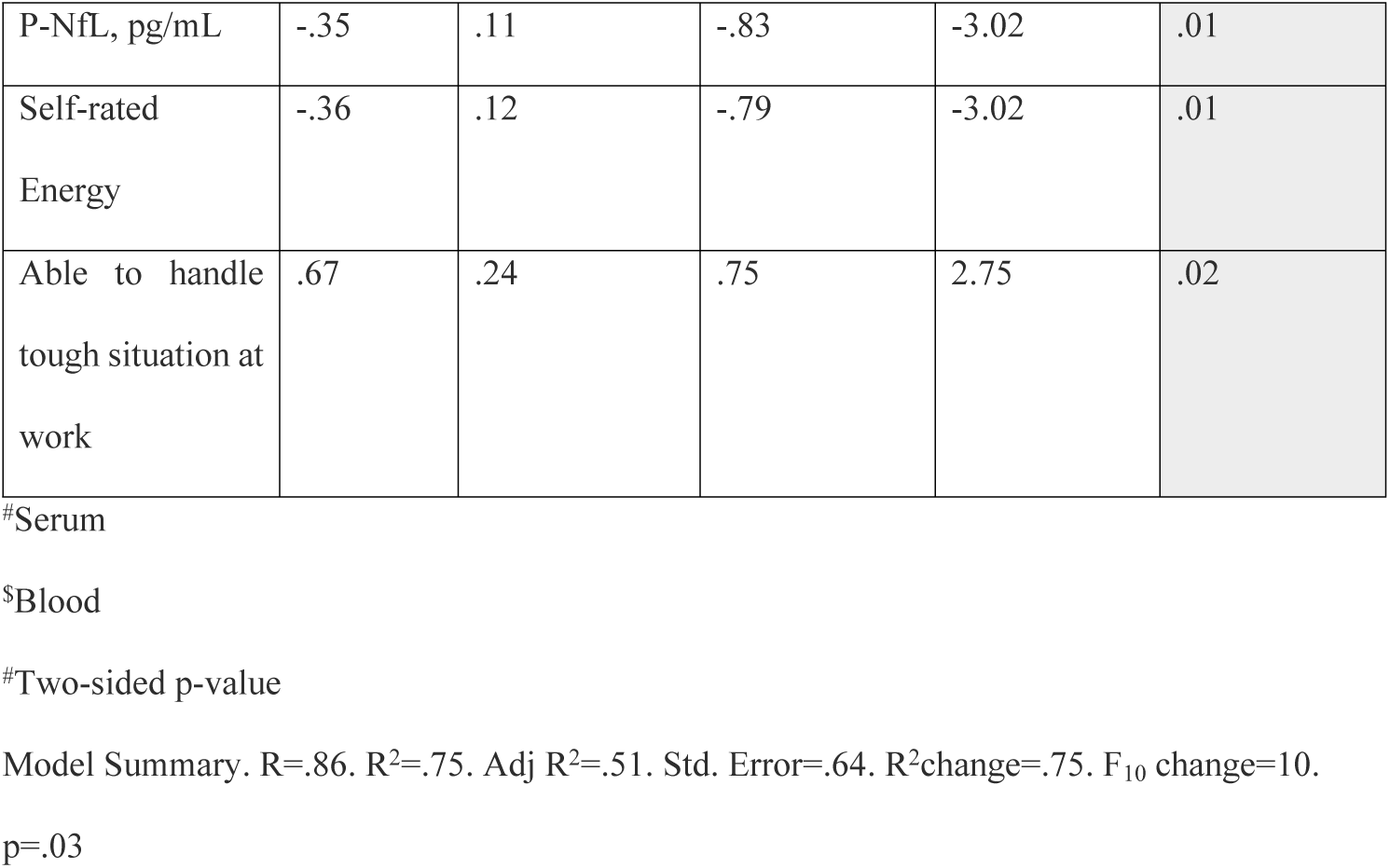
Self-rated and biomarker predictors of burnout severity category.

| Independent<br>Variable | Unstandardized<br>Coefficients |  | Standardized<br>Coefficients<br>Beta | t | p-value^ |
| --- | --- | --- | --- | --- | --- |
|  | B | Std. Error |  |  |  |
| Constant | 1.59 | 1.83 |  | .87 | .40 |
| S<sup>#</sup>-DHEA-s,<br>mg/mL | -.99 | .30 | -.73 | -3.32 | .007 |
| S-Cortisol,<br>ug/mL | -.08 | .05 | -.36 | -1.68 | .12 |
| S-IL-10, pg/mL | .03 | .02 | .34 | 1.50 | .16 |
| S-IL-6, pg/mL | .05 | .06 | .16 | .83 | .42 |
| S-IL-8, pg/mL | .63 | .30 | .48 | 2.12 | .06 |
| P<sup>s</sup>-BDNF,<br>pg/mL | .001 | .001 | .13 | .68 | .51 |
| P-NGF, pg/mL | -.01 | .04 | -.03 | -.15 | .88 |
| P-NfL, pg/mL | -.35 | .11 | -.83 | -3.02 | .01 |
| Self-rated<br>Energy | -.36 | .12 | -.79 | -3.02 | .01 |
| Able to handle<br>tough situation at<br>work | .67 | .24 | .75 | 2.75 | .02 |

#Serum

Plasma concentration of the hypothesized neuronal strain marker NfL was inversely associated with serum concentration of neuronal toxic stress biomarker cortisol (r=-.21, p=.04).

There was no association between the two measures of cognitive impairment (1. Ability to concentrate, and 2. Lapses in memory at work) and blood biomarkers reflective of the four physiological systems. Moreover, there were no associations between self-reports associated with burnout (Energy, Stress, Ability to cope with a challenging situation at work, Ability to concentrate, Lapses in memory at work) and any of the biomarkers reflective of the four physiological stress adaptive systems.

The likelihood of scoring at or above the median (Table 3 lists the median values) for participants in burnout category 1 (no-burnout) was cross-tabbed versus the four categories of burnout. There were significant associations between whether one was below or equal and above the median for energy (p=.002), ability to concentrate (p=.002), stress at work (p=.002), and NfL (p=.009). The sum scores of whether a participant scored below (score 1) or equal to and above (2) the median was created. Scoring was reversed when warranted to ensure higher index scores reflected a less desirable outcome. There was no association between the index score and the four burnout categories.

Finally, we analyzed whether there was any association between typically working day- vs night shift and self-rated and biological variables discussed above. The only significant difference was that the concentration of NfL in plasma was significantly higher in those typically worked day shifts (mean 5.71 (S.D. 2.75) pg/mL) versus night shifts (3.57 (1.65; F_1_ = 8.97, p = .003.)

## Discussion

Results show that one in every three nurses reported experiencing significant symptoms of burnout. These rates are lower than those based on converting the frequency of symptoms associated with burnout to a dichotomized outcome of burnout vs. no burnout (3, 16, 18].

However, rates reported here are based on the nurses’ perception of burnout rather than estimating the prevalence of burnout based on symptoms that lack rigorous validation in terms of cut-offs for identifying true burnout vs none.

In support of hypothesis 1, findings show a clear association across the categories of burnout and decreased self-rated energy, ability to concentrate, and work-related stress.

However, burnout was not associated with lapses in memory at work. Despite these physiological and cognitive effects, the absence of a direct relationship between burnout and memory lapses may be attributed to compensatory and energy-draining mechanisms resulting in increased cognitive load, energy consumption and, eventually, an accelerated burnout process. Task switching and multi-tasking, common in nurses actively working with electronic health records (EHR), are associated with both errors and near errors [63]. Sub-optimally designed EHR, common in clinical practice, results in increased provider cognitive load and worsened performance [64]. Further research is clearly needed to better understand neurobiological mechanisms linking suboptimal EHR design and task interruptions. For example, how do built-in redundancies such as electronic health records alerts, team-based safety checks, and standardized clinical protocol that reduce reliance on short-term recall change cognitive load, energy consumption, and ultimately burnout [63]. These findings suggest that while burnout impacts energy levels and concentration, cognitive scaffolding in the work environment may buffer against noticeable memory deficits.

Regarding hypothesis 2, participants in the combined two highest categories of burnout, Cat 4 and 5, had elevated levels of serum IL-10 vs the lowest burnout category. Increased levels of IL-10 may be the body’s adaptive mechanism to fight fatigue and low energy in persons with severe burnout since IL-10 is pro-energetic and anti-inflammatory [20, 37)] Moreover, since IL- 10 is anti-inflammatory, increased circulating levels of IL-10 might signal an attempt of the body to fight system inflammation, possibly associated with burnout [20, 37]. Work-related fatigue is closely related to low-energy syndrome, one of the most common signs of burnout. Prior work has related low-energy syndrome to lower levels of the anti-inflammatory marker interleukin-10 [20, 37].

Another novel finding is that burnout is associated with decreased serum levels of the anabolic and restorative hormone DHEA-s. This suggests that the body’s restorative capacity is reduced in patients with burnout. In addition, there is an inverse association between burnout and plasma levels of NfL, at higher concentration a marker of neuronal damage, and, if severely elevated, risk of Alzheimer’s disease and related dementias [65]. These findings suggest a need to study in further detail how burnout might be related to neuronal-axis strain and injuries and brain stress [46]. Interestingly, this adaptive response may be independent of changes in neurotrophic processes since plasma levels of NGF and BDNF were not associated with burnout. Based on these findings, it is warranted to study whether NfL at levels found in participants not diagnosed with Alzheimer’s disease is reflective of burnout severity. Cortisol is known to cause neuronal axis injuries and stress induces increased section of cortisol [66]. The inverse association between blood concentrations of cortisol and NfL might suggest that a lower concentration than what is seen in at-risk Alzheimer patients, NfL might be a marker of neuronal growth rather than injuries. Prior work has reported an association between symptoms of burnout in healthy individuals and lower levels of BDNF [4, 47, 49]. However, to our knowledge, no previous study has applied an integrated model where both self-reports of individual health and work-related cognitive impairments and biomarkers reflective of key neuroanatomical and physiological systems purported to be involved in the burnout are compared vs a validated measure of burnout.

Regarding hypothesis 3, it was unexpected that the ability to cope with challenges at work was associated with higher burnout. It might reflect that nurses that feel that they are dealing with tough situations at work do so at the cost of excessive energy expenditure resulting in the depletion of energy. Coping is also aimed at preserving status quo. The “energetic cost of allostasis and allostatic load” requires the body to spend additional energy to maintain stability in the face of stressors [67].

The construct of burnout continues to be challenging in terms of theoretical framework, definition, measurement, mechanisms, and risk and protective factors [3, 16]. Consequently, measuring and effectively addressing burnout in the clinical setting has been challenging. There is a void of validated theories and effective treatment, and burnout threatens the health and well- being of providers and patients for whom they care [37, 68]. Most instruments used to assess burnout are too lengthy for a high response rate among busy providers. The most common instruments have also been challenged from a psychometric perspective [3, 69]. The consequences of both psychometric and feasibility concerns are that the prevalence estimates of burnout are imprecise and unreliable [3]. In the current study, we used the single-item, validated burnout measure developed by Rohland et al. [61]. Moreover, in addition to measuring nurse participants’ degree of burnout, ratings were related to personal well-being, e.g., self-rated health and energy and, importantly, biomarkers reflective of neuroendocrine and cognitive systems closely associated with the construct of burnout. Rather than merely asking participants about their feelings of burnout, we suggest that, when possible, relevant biomarkers are added to the assessment. Biomarkers provide an opportunity to expand the studies of burnout to also include plausible biological disease mechanisms. Furthermore, as suggested by our findings of an association between burnout severity and self-perceived coping, merely assessing a person’s recognized emotions might not fully capture the entire width of such a complex process as burnout.

There are several limitations to the study that should be recognized. Although the study included a rather large number of nurses that provided blood samples, the study was cross- sectional and done at one large midwestern teaching hospital. This makes it impossible to determine the cause-and-effect relationship. Although the design aimed at having blood samples drawn as close as possible following the nurses had had responded to the survey, these two events happened a few weeks apart. Thus, self-reported well-being and work characteristics might have changed at the time blood was drawn. Prior work has shown that changes in work conditions at the aggregate level typically take time to occur and several of the biomarkers we choose are reflective of long-term processes that don’t changes markedly over a shorter time period [44, 70–72]. We also lack independent measures of work characteristics and information of work conditions and coping style is all based on self-reports.

In conclusion, the current study finds an association between burnout, low energy, the ability to concentrate, and lower levels of the restorative hormone DHEA-s and the neuro-axonal protein NfL. Findings of decreased NfL concentration, traditionally considered a marker of neuronal strain, in burned-out nurses warrant further studies. It’s intriguing to speculate that, since there was an inverse association between blood concentrations of NfL and the catabolic and neuro adverse stress hormone cortisol, NfL in persons still in the workforce might be an indicator of neuroplastic rather than neurostrain processes. Severity of burnout is associated with increased concentration of IL-10, an interleukin that is both pro-energetic and anti-inflammatory. This suggests that there is an active physiological process aimed at counteracting decreased energy in burned-out nurses. The increased concentration of IL-10 in burned-out participants warrants further studies of the state of systemic inflammation in burnout. The finding that decreased ability to cope with challenges at work is associated with higher burnout also requires further study.

The results also point to the need for inter-disciplinary prospective studies to delineate the temporal association between burnout and key biomarkers of neuroplasticity and neuronal strain. Burnout research, and clinical practice, would likely benefit from adding biomarkers when assessing burned-out persons since biomarkers complement the understanding of burnout. Enhanced understanding of psycho-biological disease processes contributing to burnout would also contribute to more effective strategies to prevent and treat the burnout epidemic.

## Author Contribution

Bengt B. Arnetz was the academic principal investigator, responsible for overseeing all Michigan State University-based tasks, and responsible for writing – the original draft preparation.

Bengt B. Arnetz, Judy Arnetz and John vanSchagen were responsible for funding acquisition, supervision, and validation.

Bengt B. Arnetz, Jackeline Iseler, Michelle Pena, Nicolina Evola, John vanSchagen, and Judy Arnetz were responsible for conceptualization, data curation, investigation, methodology and creation of models, project administration, visualization, writing - review and editing.

Bengt B. Arnetz and Judy Arnetz are responsible for formal analysis.

Eamonn Arble is responsible for providing specific analytical resources and analysis as well as writing - review & editing.

Scott E. Counts and John S. Beck were responsible for procuring and overseeing all biomarker analyses, including validity checks. They were also engaged in writing – review & editing. Dr. Counts was responsible for all supervision of biomarker analytical methodologies and execution.

## Data availability

The anonymized data set will be shared upon reasonable request to the corresponding author.

### Acknowledgements.

We are very grateful to all the nurses who, despite a busy schedule, volunteered to take part in the study.

